# Instruments to assess Evidence-Based Practice among healthcare professionals: a systematic review

**DOI:** 10.1101/2021.08.25.457703

**Authors:** Anderson Martins da Silva, Daniela Pereira Valentim, Adriana Leite Martins, Rosimeire Simprini Padula

## Abstract

The study makes it possible to select the most appropriate instruments to evaluate the use of Evidence-Based Practice (EBP) among health professionals. The objective of this study was to assess the measurement properties, summarize and describe the instruments that evaluate the use of EBP in health professionals, currently available through the update of the systematic review. The study was conducted and reported according to recommendations of the PRISMA checklist. A systematic search was conducted in the databases: PubMed, Embase, CINAHL and ERIC. In addition, three groups of search terms: EBP terms; evaluation; cross-cultural adaptation and measurement proprieties. They included studies that showed assessment tools of EBP in healthcare workers in general publication of full-text scientific articles, which tested the measurement properties and publication of an article in English. Searches included published studies from 2006 until July 2020. Evaluation of the methodological quality of the studies was conducted according to the COSMIN initiative. 92 studies were included. Forty new instruments have been identified to assess EBP. From these, most were developed for nursing professionals and physiotherapists. More than 48% of studies have American and Australian English as their native language. Only 28% of the studies included students in the samples. Reliability was considered appropriate (sufficient) in 76% of the instruments. The COSMIN checklist classified 7 (seven) instruments as being suitable for use in the target audience. However, Fresno Test remains the most appropriate instrument for assessing the use of EBP in healthcare professionals. 40 new instruments that assess EBP have been identified. Most are consistent and reliable for measuring the use of EBP in healthcare professionals. The *Fresno Test,* in a list of seven reliable and valid instruments for analysis, remains the most used and the one that most assesses the domains of EBP.

## 1. Introduction

For more than three decades, Evidence-Based Practice (EBP) has been considered an essential competence to guide decisions in clinical practice ^1–4^. This movement is increasingly consolidated among health professionals who believe in the value of research evidence to guide their clinical decisions ^5, 6^. The Evidence-Based Practice (EBP) can be defined as the conscious and judicious use of the best evidence science to guide decisions about health care ^7–11^. To this end, it incorporates knowledge related to high-quality clinical research, professional knowledge and patient preferences ^6, 12–14^. However, the EBP is important for patients, professionals and health services as it provides safe and effective interventions, better diagnosis and prognosis of the patient and reduces health costs ^15^.

Several strategies have been described to assess the use of this practice, among which, the use of measurement instruments^16^. Among the hundreds of instruments available ^4,17–19^ the majority are self-explanatory questionnaires based on clinical scenarios. A systematic review published in 2006^17^, identified more than a hundred instruments that assess the use of EBP in health professionals. The study pointed out that most of the instruments were administered to medical professionals and students ^4, 17^. Among EBP skills, the acquisition and evaluation of scientific evidence were the most commonly evaluated. The test of at least one type of validity was demonstrated in 53% of the instruments, however, only 10% established three or more types of validity. In addition, instruments were identified with objective measures to assess behaviors individually, as well as instruments to determine the effectiveness of EBP curricula^17^.

Seven^20–27^ of the 104 instruments identified in this review were classified as level 1 instruments, as they presented properties of reliable measures for inter-rater reliability and internal consistency, in addition to three or more types of validity tested^6^. Among which, only the Berlin Questionnaire and Fresno Test assess the stages of EBP comprehensively ^17^. Both were developed to evaluate the teaching of EBP with students and medical professionals ^20, 22^. However, the Fresno Test is the only instrument that, in addition to assessing all stages of the EBP adoption process, presents an assessment through more realistic clinical scenarios, enabling the assessment of applied competences and skills ^12^.

Systematic reviews are one of the ways to search and use these measurement instruments, as they enable the identification of reliable instruments ^4, 19, 29^. However, the methodological quality of the studies included in a review can directly interfere in the conclusion, in the estimates of the effect and on the validity of the study^30^. Although the study by Shaneyfelt et al. (2006) ^17^ is the most recent systematic review on the subject, the criteria adopted for analyzing the properties of the instruments did not follow methodological guidelines for developing a review. In the study, no guidelines were described for the analysis of the methodological quality of the included studies, as well as for the evaluation of the properties of measures extracted from each instrument. In addition, the use of a checklist to classify each measure property evaluated was not reported.

In view of the relevance of the subject in the last decade and the time elapsed since this review, it is possible that other instruments with acceptable methodological quality have emerged. To date, still no updated systematic review summarizes all the necessary information on these measurement instruments.

Therefore, it is necessary to update the systematic review of the instruments for assessing the use of EBP, to complement these aspects and provide the updating of existing tools. The study aims to evaluate the measurement properties, summarize and describe the instruments that evaluate the use of EBP in health professionals, currently available through the update of the systematic review.

## 2. Method

### Protocol and registration

This is a systematic review study on instruments for assessing the use of EBP, verification and analysis of measurement properties tests. This review was conducted and reported according to recommendations from the PRISMA checklist (Preferred Reporting Items for Systematic Reviews and Meta-Analyzes). It was registered under number CRD-42018103212 in PROSPERO (International prospective register of systematic reviews) and can be accessed at https://www.crd.york.ac.uk/prospero/#recordDetails. The study was submitted to and approved by the Research Ethical Committee Protocol of Universidade Cidade de São Paulo - UNICID No 3.636.011/2019.

### Search strategy

To identify the EBP assessment instruments, a systematic search was conducted in the databases: PubMed, Excerpta Medica database (EMBASE), Cumulative Index to Nursing and Allied Health Literature (CINAHL) and Educational Resources Information Center (ERIC).

Three groups of search terms were used: **EBP terms**: evidence-based medicine; critical appraisal; clinical epidemiology; clinical question; medical informatics; information storage; information retrieval; Attitude to Health; Health Education; Competency-Based Education; Students, Public Health; Perception. **Evaluation terms**: program evaluation; program assessment; questionnaire; scale; index; instrument; evaluate; measurement; Test. **Terms of cross-cultural adaptation and clinimetry**: Cross-Cultural Comparison; Validation Studies; Psychometrics; Reproducibility of Results; Statistics; Observer Variation; Methods; Comparative Study; Outcome Assessment; Discriminant Analysis; translation; validation; agreement; clinimetric; construct validity; concordance; internal consistency; interpretability; measurement properties; reproducibility; -responsive; reliability; measure; subscale; sensitive; responsiveness.

As search strategies, the following steps were adopted: within each group there was a combination of terms with the OR particle. Subsequently, the groups were combined with the AND particle. In addition, manual searches were conducted carried out newspapers and periodicals on education and EBP, specific search by authors and contact with researchers, In addition to internet sites. Studies that describe instruments for assessing the use of EBP in students and health professionals were identified. These studies were analyzed for their methodological quality and their instruments were described and differentiated as to the scope of the EBP domains covered and the properties of measures tested.

The studies selected were the ones that: (1) presented an evaluation instrument; (2) evaluated EBP in health professionals in general; (3) presented the publication of a full-text scientific article; (4) tested the measurement properties and (5) publication of an article in English. Studies that (1) were published in another language were excluded; (2) evaluated teaching, but not evidence-based practice.

Searches were limited by publication date and included studies published from 2006 to July 2020. The searches were performed again before the final analysis on July 30^th^ 2020 and other studies could be retrieved for inclusion. The results of the search strategies were imported into the ENDNOTE® X9 software and the duplicates were removed. The deadline for the publication of a study with exclusion criteria was May 2006.

### Criteria for eligibility of studies

The inclusion criteria for searches for EBP assessment instruments were 1) to quote in their title or in their summary of the descriptors; 2) that the instrument or strategy for the use of EBP has assessed the patient’s skills, attitudes, behaviors or results; 3) that the instrument or strategy had a sufficient description to allow an analysis; 4) that the instrument should present results of measurement proprieties tests.

To standardize the concepts and terminologies used in this study, the BOX of definitions of variables and terminology described by Shaneyfelt et al. (2006) ^17^ was considered. The domains for EBP were defined as knowledge, skill, attitude, behavior and viability. The skill domain considered the participants’ ability to: ask, acquire, evaluate and apply the evidence in clinical decision-making. The behavior domain considered the performance of the participants for performing EBP steps in practice; execution of evidence-based clinical maneuvers and; achieve favorable results with the patient.

### Evaluation of the methodological quality of eligible studies

The data regarding the measurement properties were extracted from each study and analyzed according to the the COSMIN (COnsensus-based Standards for the selection of health Measurement INstruments) initiative ^34–36^. The systematic assessment of the methodological quality of each instrument considered three quality domains: reproducibility; validity and responsiveness. Each domain contains one or more measurement properties. The reproducibility domain considered three measurement properties: internal consistency; reliability and measurement error. The validity domain also considered three measurement properties: content validity; construct validity and criterion validity. The responsiveness domain considered only one measurement property, called responsiveness. The properties of measures that contain one or more aspects were defined separately: content validity included face validity and construct validity includes structural validity, hypothesis testing and cross-cultural validity ^34, 37–40^.

To classify each measurement property, the COSMIN guidelines were used for systematic reviews of Patient Reported Outcome Measures (PROMs) ^35^. The guideline consists of 10 boxes, recommended for obtaining global scores for the methodological quality of studies in systematic reviews.^35^. The score for each item in a box was obtained considering a 4-point rating scale (V = very good; A = adequate; D = doubtful; I = inadequate) ^35^.

Subsequently, the methodological quality of the studies was classified as excellent, good, reasonable or poor, based on the scores of the items in the corresponding box. This classification took into account the worst score for a given box. Thus, a box that classified some items as excellent, however presented an item as poor, was classified as of low methodological quality^37^.

### Data extraction and analysis

After selecting the studies that met the inclusion criteria or that made it impossible to be sure that they should be excluded, an initial analysis of the titles and abstracts was conducted. Subsequently, all selected articles were obtained in full and examined according to the established inclusion criteria.

From the selected articles, data regarding the tested psychometric properties were extracted. A COSMIN Risk of Bias Checklist was used for systematic reviews of PROMs^35^. The checklist consists of nine boxes, including: Box A (development); Box B (content validity, including face validity); Box C (structural validity); Box D (internal consistency); Box E (cross-cultural validity); Box F (reliability); Box G (measurement error); Box H (criterion validity); Box I (construct validity - hypothesis tests) and; Box J (responsiveness) ^35, 37^.

Other aspects of the instruments identified were also extracted, such as the sample size, the target audience, the year of publication and the method of application of the instrument, as well as the steps and domains for adopting the EBP proposed by the instrument. The process of evaluating the methodological quality of the studies was carried out by two independent evaluators (MAS and DPV). Possible disagreements during the process were resolved by an independent reviewer (RSP).

## 3. Results

In total, 6,429 studies were found in four databases searched, among which 92 were considered eligible for data analysis. From the 92 eligible studies, 46 unique instruments were identified (appendix). No divergence of opinion was found between the evaluators and the independent reviewers for the eligibility of the included studies. The study’s PRISMA flowchart is shown in figure 1.

**Figure 1.**
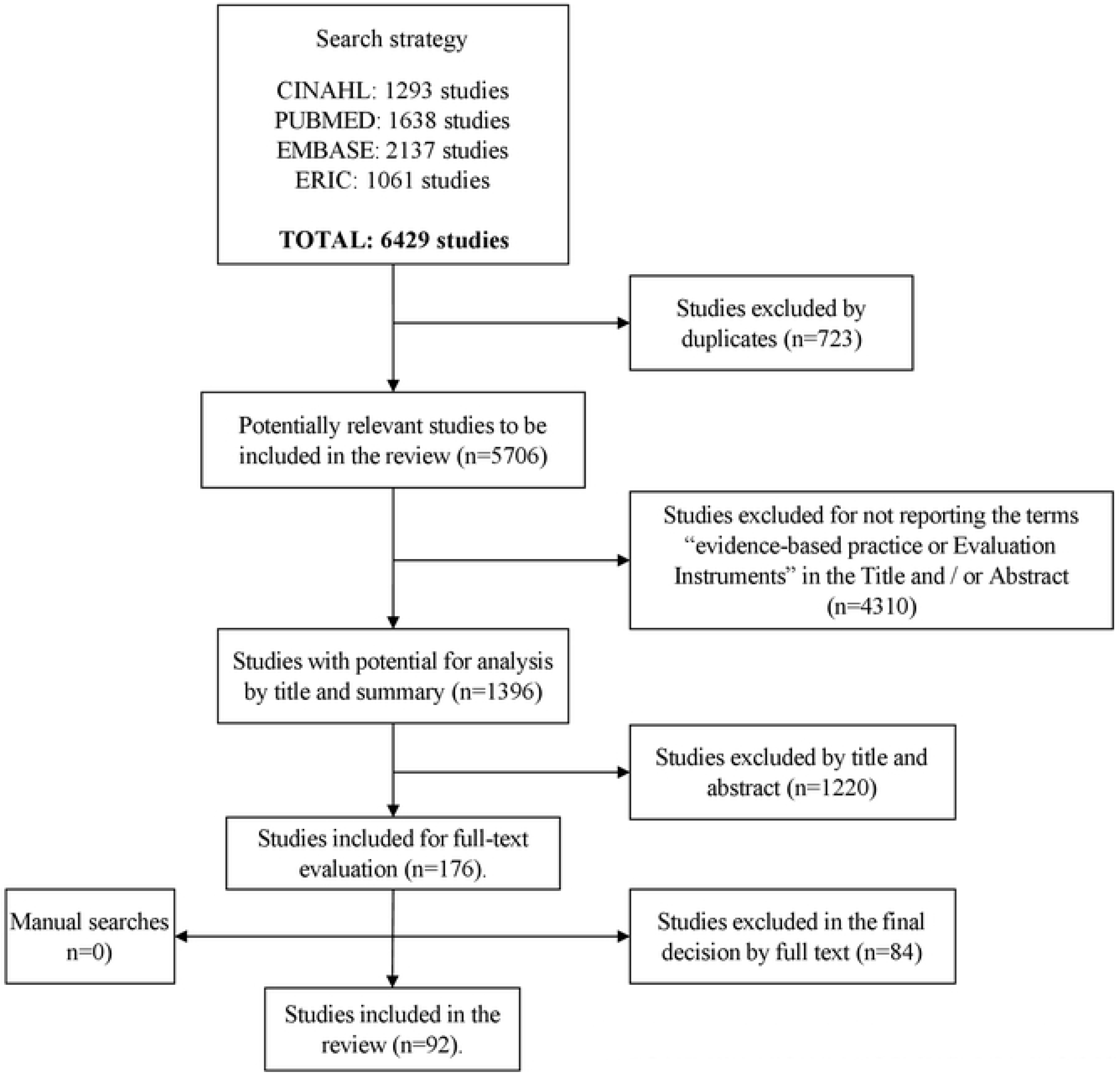
PRISMA flowchart of the study.

### Description of the studies

The general characteristics of the studies included in the review are shown in table 1. Nursing professionals were the most evaluated samples (59.8%), followed by Physiotherapists (27.1%) and Physicians (19.6%). Most of the instruments used were aimed at evaluating graduated professionals (71.8%). The USA and Australia were the countries that presented the largest number of publications (48.9%). Most publications occurred in the last 4 years with (50.0%) and with a sample (N) between 101 and 500 participants (43.5%). The name of the instrument used in 3 studies was not reported, despite the description of the characteristics of the instruments. Eleven studies reported the use of more than one instrument to assess EBP.

**Table 1.**
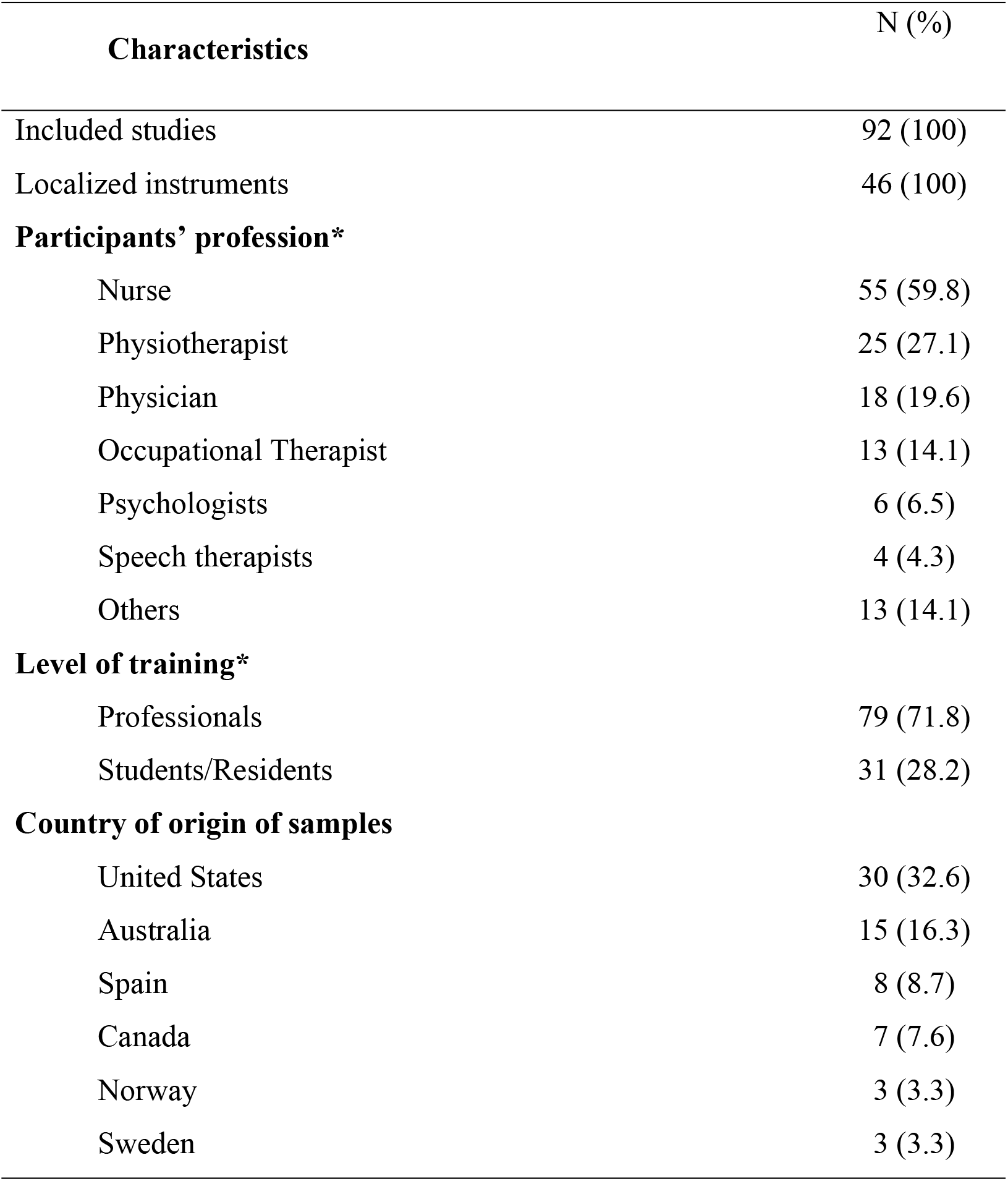
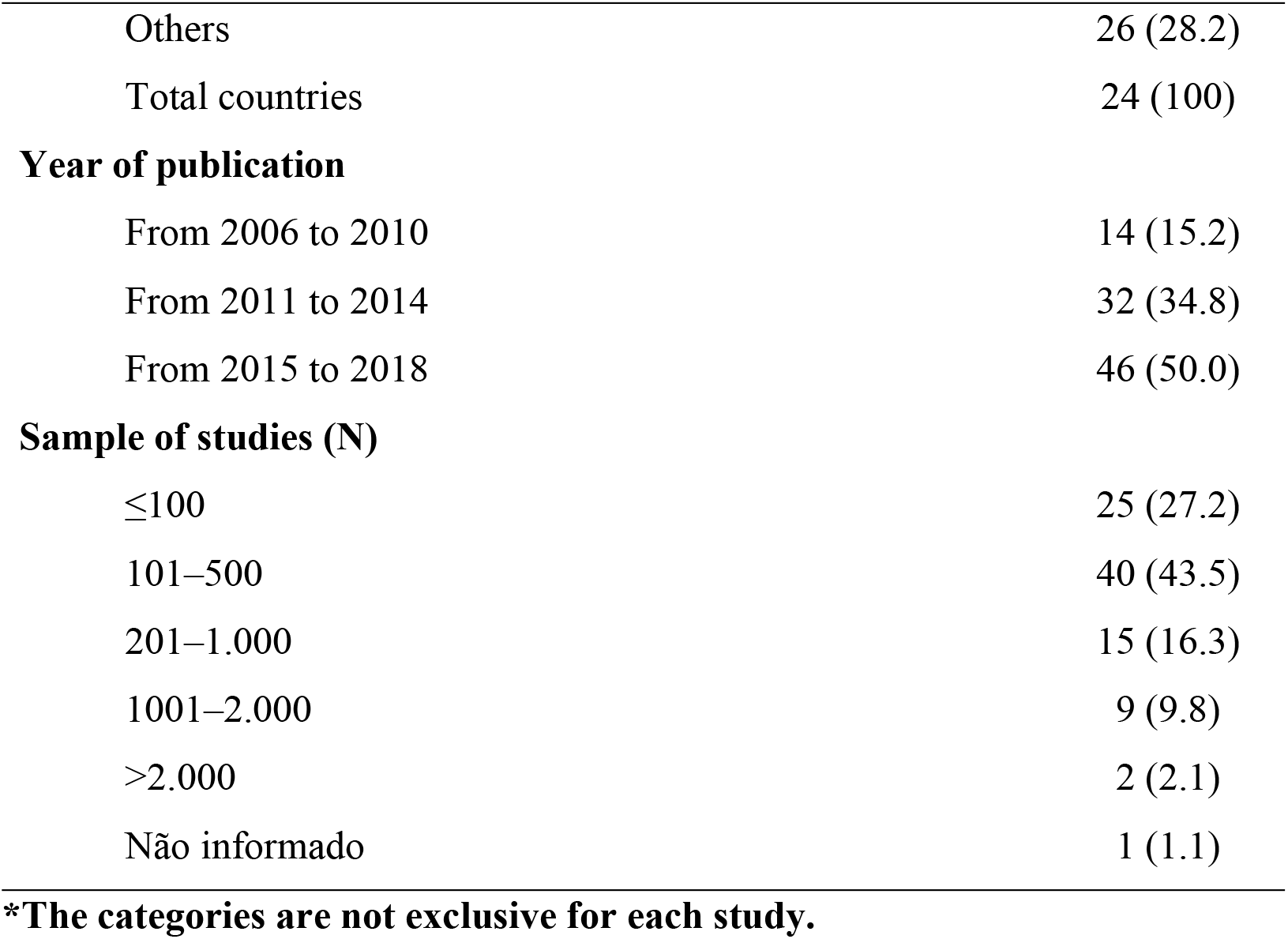
Characteristics of the included studies (N = 92)

### Description of EBP Assessment tools

As for the characteristics of the EBP assessment instruments located, the Fresno Test (FRESNO) was the most used (12.0%), followed by the Evidence-Based Practice Attitude Scale (EBPAS), (10.9%) and Nurse Manager EBP Competency Scale (8.7%), (table 2). Among the 46 instruments identified, the majority (78.2%) consisted of closed questions and a Likert scale of answers, administered to the participants in printed form (44.5%). Most of the studies had cross-sectional designs (73.9%), the objective was to develop new instruments and reduced versions (34.8%) and to conduct formative assessment (individual), (83.7%). The EBP Domain most evaluated was attitude (63.0%), followed by knowledge (55.4%) and skills (28.2%). Among the EBP skills, the participants’ ability to “apply” the evidence in clinical decision-making and the ability to “critically evaluate” this evidence for its validity and applicability represented most of the instruments. Still, 59.0% of the instruments evaluate between 1 and 2 stages of 5 described for the process of adoption of EBP.

**Table 2.**
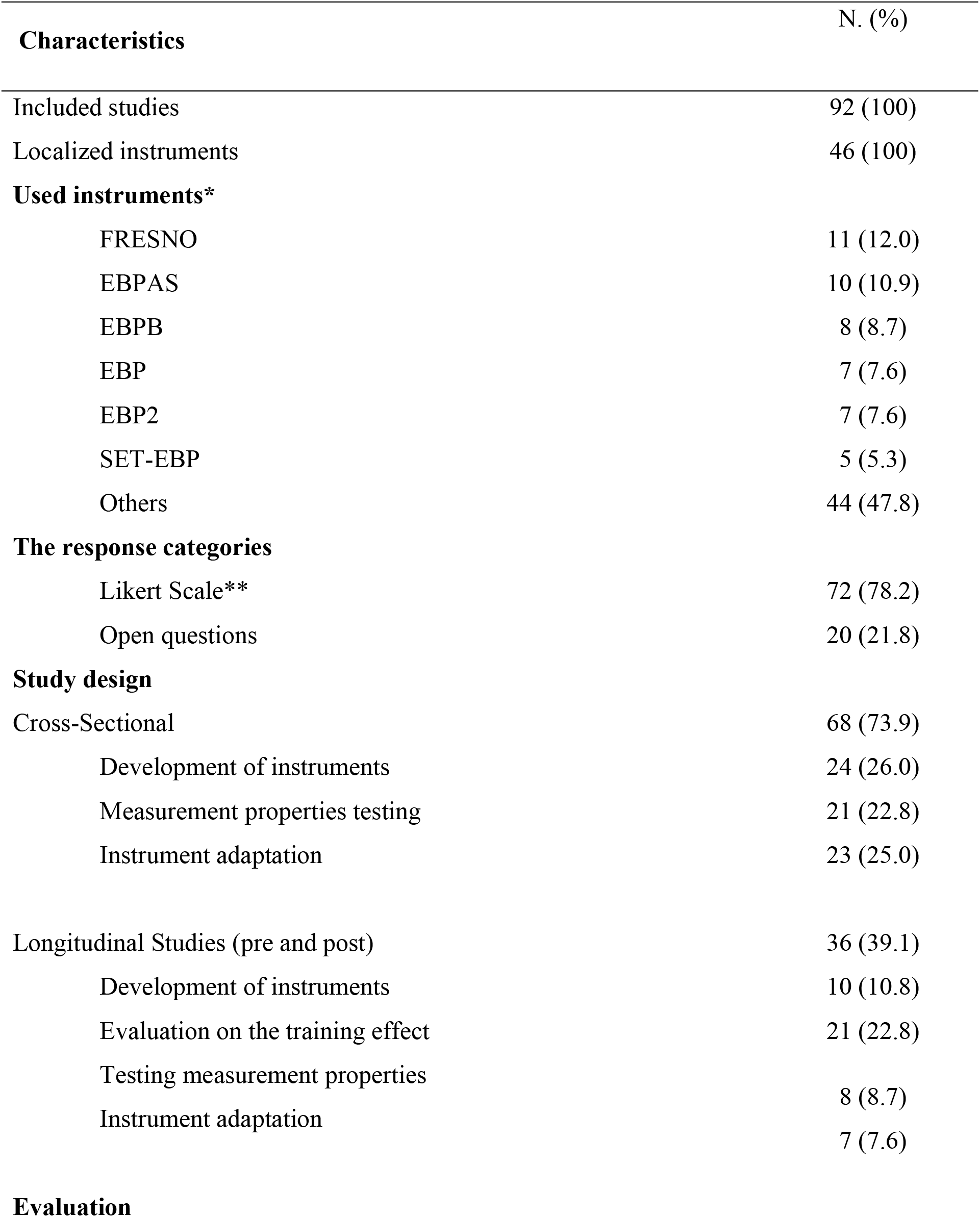
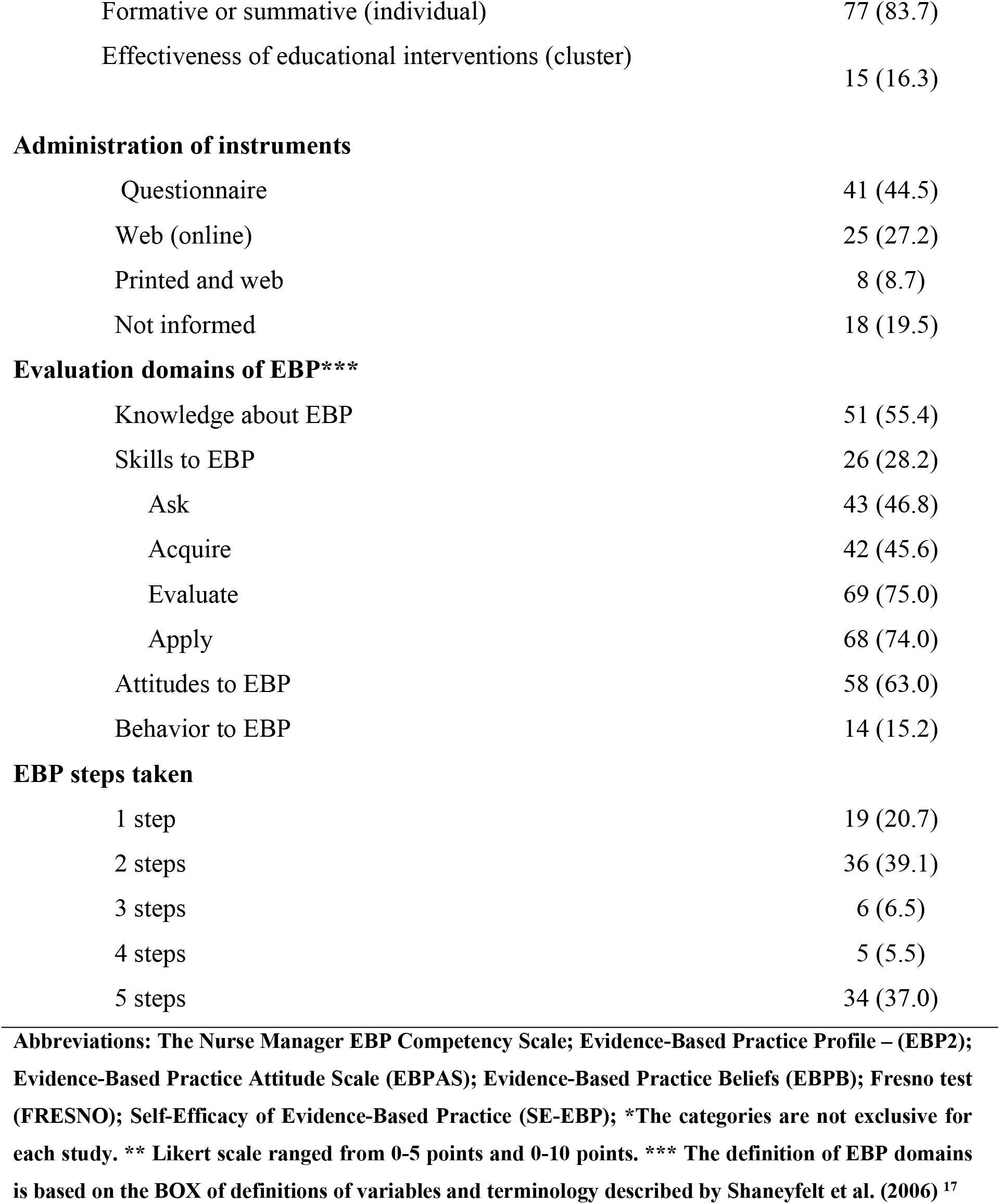
Characteristics of the EBP assessment instruments (N = 92).

The researchers performed at least 1 type of validity test on 73% of the instruments evaluated in the study (table 3). Reproducibility was tested on 90% of the instruments, using internal consistency. Responsiveness was tested on less than half of the instruments (30%). The classification for the evaluation of each tested measurement property, considered the evaluation sufficient (+), according to the COSMIN checklist. Internal consistency was the reproducibility test that presented the highest percentage of sufficient evaluation (+), with 76%. Construct validity was assessed as sufficient (+) in 67% of the instruments. The absence of risk of bias for the quality of the studies was identified in more than half (54%) of the studies. The studies that presented a “high” quality of the evidence represented 50% of the total of evaluated studies

**Table 3.** Measurement proprieties characteristics of EBP instruments (N = 92).

| **Measurement property** | **Tested**<br>**N (%)** | **Sufficient (+)\***<br>**N (%)** |
| --- | --- | --- |
| **Validity** |  |  |
| Content | 54 (58.6) | 46 (50.0) |
| Construct | 68 (73.9) | 62 (67.3) |
| **Reproducibility** |  |  |
| Reliability | 81 (88.0) | 46 (50.0) |
| Internal consistency | 83 (90.2) | 70 (76.0) |
| Measurement error | 32 (34.7) | 31 (33.6) |
| **Responsiveness** |  |  |
| Responsiveness | 28 (30.4) | 27 (29.3) |
| **Other evaluation criteria** |  |  |
| Risk of Bias | 50 (54.3) | Absent** |
| Quality of evidence | 46 (50.0) | High*** |

**General classification according to the COSMIN checklist: \* Evaluation of measurement properties**
**\*\* Risk of Bias \*\*\* Quality of Evidence.**

### Categorization of Study Quality

As in the study by Shaneyfelt et al. (2006), we defined the 7 (seven) instruments that were classified as LEVEL 1 (table 4). These instruments were considered suitable for use in the target audience. However, we consider the COSMIN initiative to define the quality of the instruments. The instruments were defined based on the following criteria: (1) no risk of bias for the methodological quality of the studies; (2) a high-quality evidence; (3) tests of reliability, validity and responsiveness classified as sufficient, and (4) scope in the domains and stages of adoption of EBP, proposed by the instrument. The criteria were adopted considering the general classification for all studies described on the instrument. Nursing professionals were included in the samples of all instruments classified as LEVEL 1, followed by physical therapists. Nutrition professionals were tested on only 1 instrument. All instruments have a self-report response format. Except for the EPIC and FRESNO instruments, all instruments have subscales.

**Table 4.**
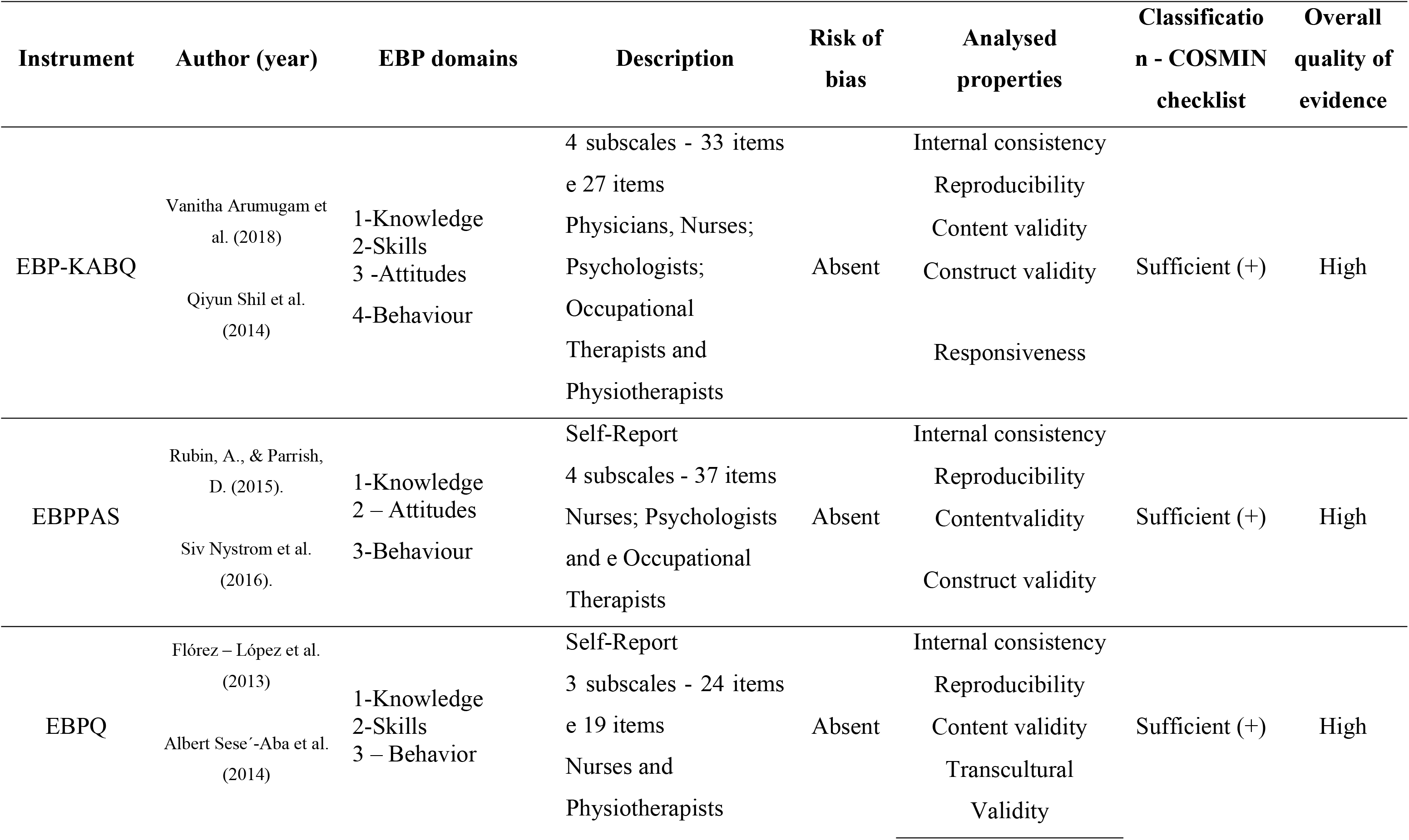
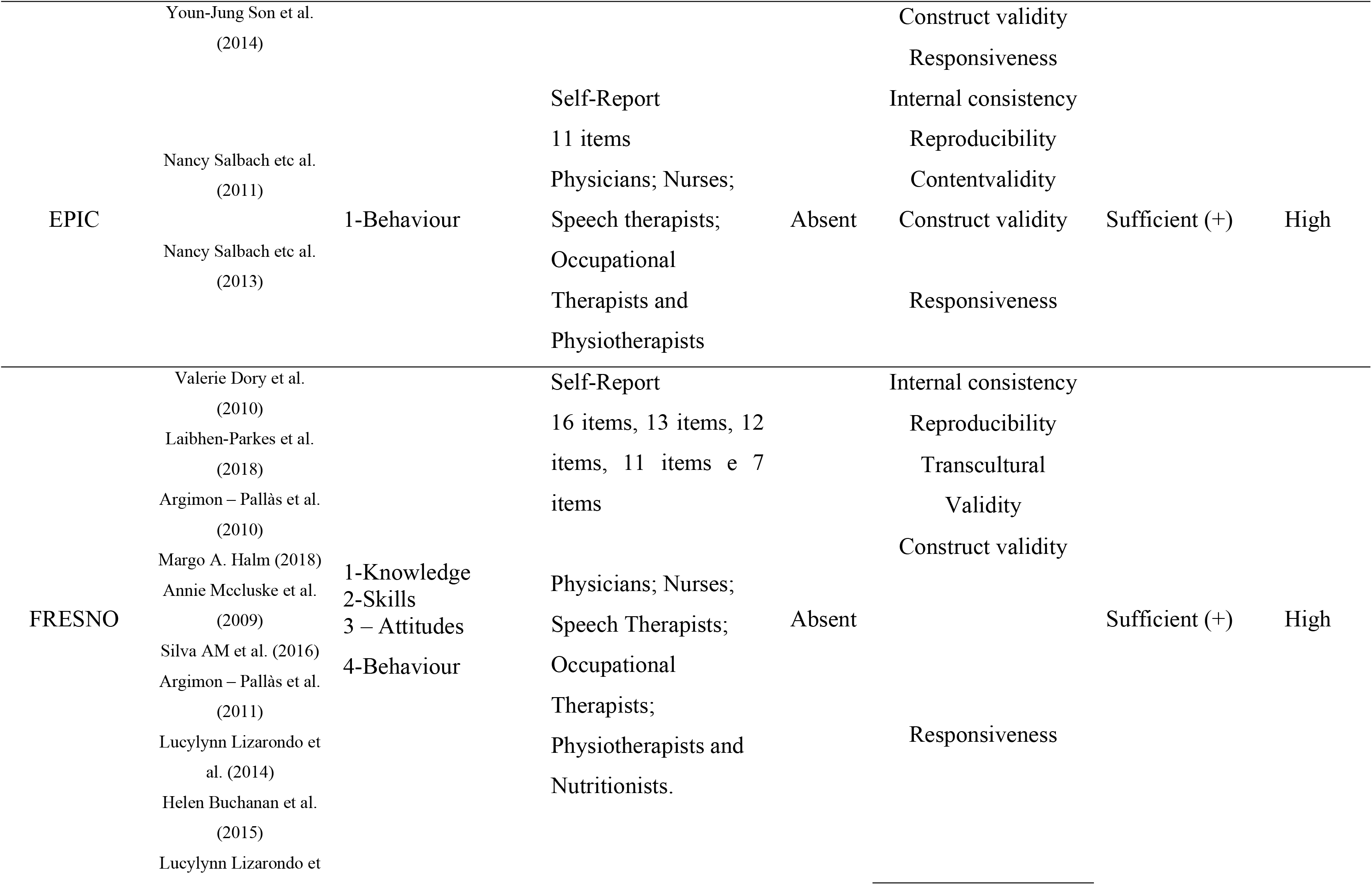
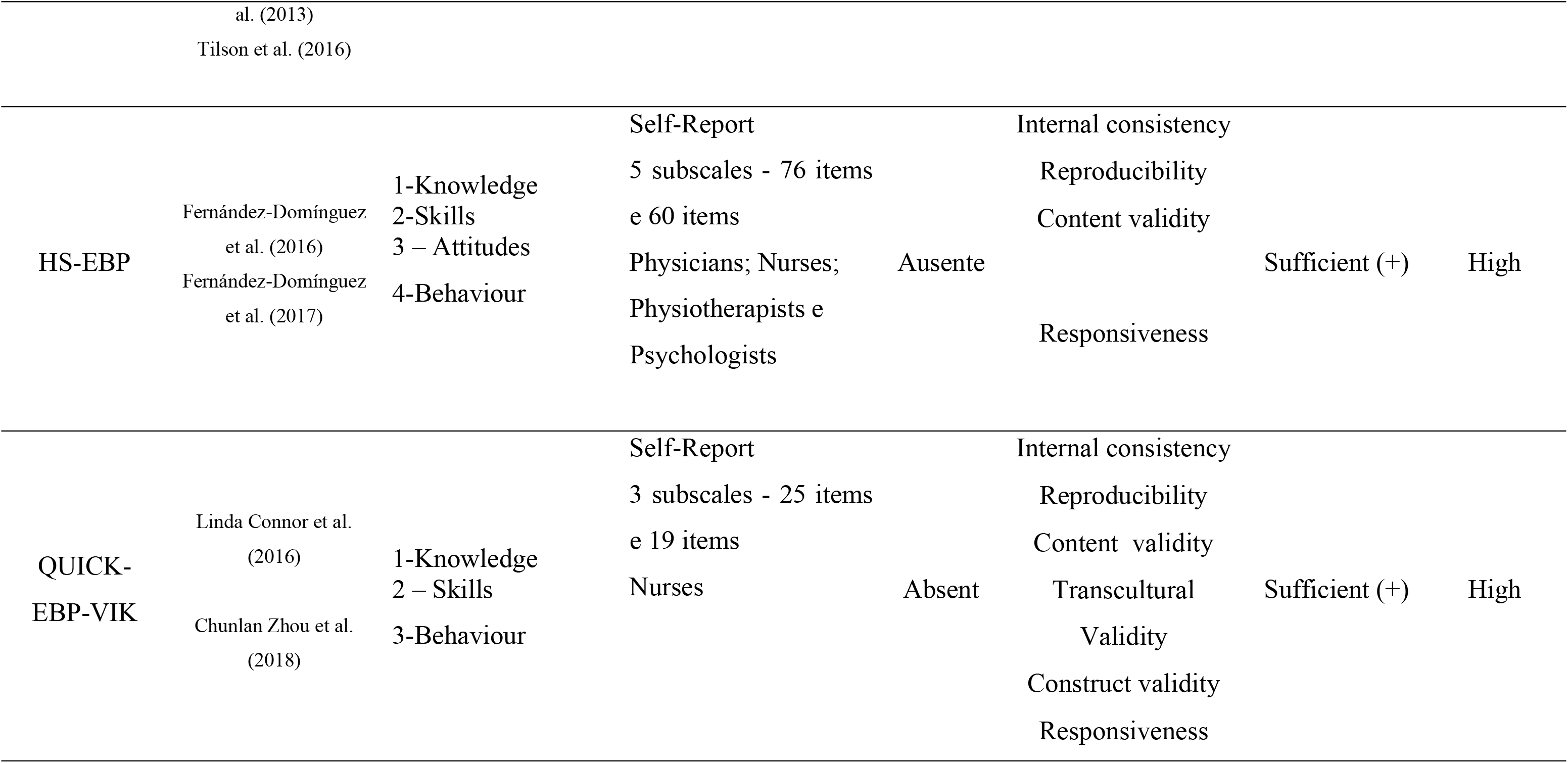
Classification of EBP assessment instruments according to the COSMIN checklist.

The EBPQ and QUICK-EBP-VIK instruments tested all the measurement properties that make up the COSMIN checklist evaluation BOX. These instruments assessed internal consistency, reliability, structural validity, cross-cultural validity, construct validity and responsiveness. However, both assess only 3 of the 4 domains proposed for the adoption of EBP. The validation studies report limitations of these instruments in relation to the response format (self-report) and sample size used. The EBPQ consists of 24 items and 3 subscales and has a short version of 19 items. The general classification of the EBPQ was based on 3 (three) studies that tested the measurement properties in nurses and physiotherapists. The QUICK-EBP-VIK consists of 25 items and 3 subscales, has a short version of 19 items and measures three domains of EBP: value (V), implementation (I) and knowledge (K). The value represents how nurses believe in the importance of EBP in their clinical practice. The implementation is related to the ability to conduct the stages of the EBP adoption process. Knowledge is related to the nurse’s execution of the EBP steps. Each question is assessed on a five-point Likert scale.

The measurement properties of the instrument were tested in 2 (two) studies with professional nurses.

Among the 7 instruments classified as NIVEL 1, HS-EBP is the instrument with the highest number of assessment items (questions), followed by EBPPAS. The HS-EBP consists of 76 items and 5 subscales and features a version of 60 items. Evaluates knowledge, skills, attitudes and behavior related to EBP in health professionals. It presents the evaluation by question using four-point Likert scales and including open fields for suggestions. Two studies tested the measurement properties of the instrument in Doctors, Nurses, Physiotherapists and Psychologists. EBPPAS was developed to assess the impact of EBP education on students and health professionals. It consists of 37 items and 4 subscales based on the five stages of the EBP adoption process. It presents the evaluation by question using 5 (five) point Likert scales. The scale makes it possible to assess the behavior of students and professionals after receiving educational intervention. It has the advantages of being relatively short, easy to use. Two studies tested the measurement properties of the instrument in Nurses; Occupational Psychologists and Therapists. As for the limitations, the high number of items in these instruments may be related to the low response rate reported in the validation studies.

EPIC is a self-report scale developed to assess the effects of EBP education among healthcare professionals. It consists of 11 items that describe the steps considered relevant to the EBP adoption process. These steps include the acquisition, evaluation and effective application of the identified evidence to solve a clinical problem. Participants are assessed for their level of confidence in the execution of each step using an eleven-point Likert scale from 0 to 11. Two studies tested the measurement properties of the instrument in health professionals. The instrument has limitations regarding the sample profile used.

The EBP-KABQ consists of 33 items and 4 subscales and features a version of 27 items. Evaluates knowledge, skills, attitudes and behavior related to EBP in health professionals. It presents the evaluation by question using six- and seven-point Likert scales. Two studies tested the measurement properties of the instrument in Doctors, Nurses; Psychologists; Occupational Therapists and Physiotherapists. However, limitations are reported in both studies related to geographic and demographic differences in the samples. In addition, the instrument is presented in English only.

FRESNO was the most used instrument by researchers. It was also the instrument that evaluated the use of EBP in the largest number of different professions among health professionals. Eleven studies evaluated EBP in Physicians; Nurses; Speech therapists; Occupational Therapists; Physiotherapists and Nutritionists.

The instrument assesses all the domains proposed for the adoption of EBP. The Fresno Test is a self-explanatory instrument where the participant must choose one of the clinical scenarios so that, from it, one can answer the open questions. It stands out for presenting the most realistic clinical scenarios, enabling the assessment of applied knowledge and skills.

## 4. Discussion

### Summary of evidence

This study was to carry out an update through a systematic review of the instruments for assessing the use of EBP and the quality of the measurement properties of the instruments. The study by Shaneyfelt et al. ^17^ carried out in 2006, it is the most recent systematic review on the subject.

In the last few decades, EBP has become an essential competence to guide decisions in clinical practice among healthcare professionals. Therefore, it is essential to constantly evaluate this practice, using reliable instruments.

Our study showed a considerable increase in new instruments aimed at non-medical professionals in relation to the study by Shaneyfelt et al. ^17^. 40 (forty) new instruments were identified. From these, most of them were developed for nursing professionals and physiotherapists. Studies with non-medical professionals included in the samples may have contributed to the emergence of new instruments. Health professionals, not doctors, were included in more than 80% of the studies. This means an increase of 67% over the 2006 study^17^.

The emergence of new instruments makes it possible to constantly assess the effectiveness of the EBP adoption process. This is an important indicator of the development and consolidation of the use of EBP among health professionals. However, even with the emergence of new studies and instruments, there are still professionals not included in this process of evaluating the use of EBP. Only two instruments were identified to assess the use of EBP with dental professionals. One of them did not present a denomination. The other is called Evidence-Based Practice Knowledge, Attitudes, Access, and Confidence Evaluation (KACE). Both were classified as “low-quality of evidence” by the COSMIN checklist. The use of EBP by Nutritionists was assessed by the Modified Fresno Test, the only instrument adapted for these professionals, presented “high-quality evidence.” Still, no instruments were identified to assess the use of EBP in Pharmacy and Biomedicine professionals.

The fact that almost half of the studies (48.9%) have American and Australian English as their native language is also a problem. Due to the fact that, although certain professions have several instruments available, few have been adapted for certain languages and countries. Thus, although health professionals increasingly believe in the value of research evidence to guide their decisions, there is a need to develop or adapt to new instruments with suitable measurement properties for use in these populations.

We evidence that in recent years, health students have been poorly evaluated for the use of EBP. Only 28% of the studies included students in the samples. These findings can hinder the development and consolidation of this practice among health professionals. The teaching of EBP and the constant assessment of competences and skills acquired by professionals during their training can alleviate the difficulty in seeking, interpreting and translating the evidence into clinical practice. It is the biggest obstacle faced by health professionals for adopting EBP. The attitude toward EBP, characterizes a skill that can also alleviate these difficulties. The attitude was the most evaluated EBP domain among studies (63%). The COSMIN checklist considered the construct validity of more than half of the instruments (67%) appropriate (sufficient). These values are higher than the study by Shaneyfelt et al. in 2006, where only 53% of the 104 EBP assessment instruments was considered appropriate for tested validity. The structural validity was appropriate for 50% of the tested instruments. Most used confirmatory factor analysis (CFA) or comparative adjustment index (CFI). However, cross-cultural validity was appropriate for only 22% of the instruments tested through the difference between groups or by the functioning of the differential item (DIF). This means that most of the instruments identified in this review are appropriate for measuring the construct they propose, for which they were developed, however, they present weakness in the tests of cross-cultural validity. This condition can make it difficult to use these instruments in certain countries or cultures. The fact that the constructs evaluated by the instruments for evaluating the use of EBP in health professionals do not have a gold standard, it was not possible to assess the criterion validity of the instruments. We only considered; the validity correlated the scores with another tool that measures the same construct (construct validity).

The reliability considered appropriate (sufficient) in 76% of the instruments allows us to affirm that most identified instruments are consistent and reliable for measuring the use of EBP in health professionals. The evaluation considered an ICC or Kappa weighted correlation coefficient ≥ 0.70 for the classification. However, many studies have used other variables to assess the reliability of the instruments. The results may justify the identification of the risk of bias for the properties of the tested measures and the moderate or low overall quality of the evidence in 50% of the studies included in this review.

The COSMIN checklist used in our study also considered the responsiveness of the instruments. However, less than half of the instruments tested (29%), were classified as appropriate (sufficient) by the COSMIN checklist. These findings show a weakness in the assessment of responsiveness of EBP assessment instruments in health professionals. Little responsive instruments are unable to detect changes in a test over time. This inability is decisive for the quality of a measuring instrument. Furthermore, it is possible that the risk of bias, present in 46% of the studies, is not related to the size of the samples but to the properties of the tested measures.

We used the COSMIN checklist to assess the methodological quality of the included studies and to classify each measure property evaluated. This ensured a systematic and transparent collection and evaluation of information, as well as an overview of the measurement properties of the instruments. It also made it possible to recommend the most appropriate instruments to assess the use of EBP in health professionals. These guidelines for the analysis of the methodological quality of the studies had not been used in the study by Shaneyfelt et al. ^17^. This analysis was defined using a form developed by the authors.

The 7 (seven) instruments classified as LEVEL 1 can be considered adequate instruments to be used in the target audience. However, even with the emergence of new instruments, we emphasize that Fresno Test 12 remains the most appropriate instrument to assess the use of EBP in health professionals. The test 12 was developed to assess the use of EBP in residents and doctors. It was described by Shaneyfelt et al. in 2006, as a reliable tool to assess all five stages of EBP objectively ^13–14^. It has been adapted for other languages ^7–15–16^ and different health professionals ^11–17–19^. This instrument can be an alternative to evaluate the use of EBP in the professions less evident in our study.

Thus, this systematic review makes it possible to select the most appropriate instruments to be used for a given purpose and supported by evidence of good measurement properties. It can also identify gaps in knowledge about the instruments available to assess the use of EBP in this audience. This can be used to design new studies on these instruments.

### Limitations

There are limitations that must be considered when interpreting the results of this review. It is possible that we have not been able to identify some assessment instruments, due to the variability of terms used in the literature related to Evidence-Based Practice. However, we searched several databases, including those that contain unpublished studies. Also, because the search was limited to the English language, other relevant studies from other non-English languages in countries may have been missed. This can introduce publication bias if such instruments systematically differ from those that appear in English Journals. As well as the exclusion of studies for not reporting the terms "evidence-based practice or assessment instruments" in the title and/or abstract, this may also have biased our analysis. Finally, although the use of the COSMIN checklist to assess the methodological quality of the included studies and to classify each measurement property evaluated contributes to the low risk of bias in our analysis, it is possible that the characteristics of some EBP assessment instruments may misclassified, particularly in determining the validity of evidence based on its relationship to other variables.

## 5. Conclusion

The update of the systematic review of the instruments for assessing the use of EBP identified 40 (forty) new instruments. Most of them are consistent and reliable for measuring the use of EBP in healthcare professionals. However, they have limitations for certain health professions. The COSMIN checklist provided an overview of the measurement properties of the instruments and classified 7 (seven) instruments as being suitable for use in the target audience. FRESNO was the most used instrument by researchers and the one that evaluates the largest number of domains of EBP, for different health professionals and in different languages. The study contributes to the development and consolidation of Evidence-Based Practice among health professionals.

## Acknowledgements

Not applicable

## Abreviations

EBP: Evidence-Based Practice
PRISMA: Preferred Reporting Items for Systematic Reviews and Meta-Analyzes
Medline: Medical Literature Analysis and Retrievel System Online
EMBAS: Excerpta Medica dataBASE
CINAHL: Cumulative Index to Nursing and Allied Health Literature
ERIC: Educational Resources Information Center
COSMIN: COnsensus-based Standards for the selection of health Measurement Instruments
PROMs: Patient Reported Outcome Measures
FRESNO: Fresno Test
EBPAS: Evidence-Based Practice Attitude Scale
EBP2: Evidence-Based Practice Profile
EBPB: Evidence-Based Practice Beliefs
SE-EBP: Self-Efficacy of Evidence-Based Practice
EBPQ: Evidence-Based Nursing Questionnaire
QUICK-EBP-VIK: Value, Implementation, and Knowledge of Evidence-Based Practice
HS-EBP: Health Sciences–Evidence-Based Practice
EBPPAS: Evidence-Based Practice Process Assessment Scale
EPIC: Evidence-based practice confidence
EBP-KABQ: Evidence-Based Practice knowledge, attitude, behavior questionnaire
KACE: Knowledge, Attitudes, Access, and Confidence Evaluation

